# RNA binding domains of heterologous viral proteins substituted for basic residues in the RSV Gag NC domain restore specific packaging of genomic RNA

**DOI:** 10.1101/2020.03.12.989541

**Authors:** Breanna L. Rice, Timothy L. Lochmann, Leslie J. Parent

## Abstract

The Rous sarcoma virus Gag polyprotein transiently traffics through the nucleus, which is required for efficient incorporation of the viral genomic RNA (gRNA) into virus particles. Packaging of gRNA is mediated by two zinc knuckles and basic residues located in the nucleocapsid (NC) domain in Gag. To further examine the role of basic residues located downstream of the zinc knuckles in gRNA encapsidation, we used a gain-of-function approach. We replaced a basic residue cluster essential for gRNA packaging with heterologous basic residue motif (BR) with RNA binding activity from either the HIV-1 Rev protein (Rev BR) or the HSV ICP27 protein (ICP27 BR). Compared to wild type Gag, the mutant ICP27 BR and Rev BR Gag proteins were much more strongly localized to the nucleus and released significantly lower levels of virus particles. Surprisingly, both the ICP27 BR and Rev BR mutants packaged normal levels of gRNA per virus particle when examined in the context of a proviral vector, yet both mutants were noninfectious. These results support the hypothesis that basic residues located in the C-terminal region of NC are required for selective gRNA packaging, potentially via initial electrostatic interactions that facilitate specific binding.

## 1. Introduction

The retroviral Gag protein directs the assembly of new virus particles at the plasma membrane, specifically selecting the viral genomic RNA (gRNA) from the milieu of cellular and viral RNAs. The mechanism by which selective packaging occurs within the cell remains incompletely understood. The Gag NC domain is essential for encapsidation of gRNA and also plays an important role in the subcellular trafficking of the Gag protein [1–6] RSV Gag is initially synthesized in the cytoplasm and then undergoes transient nuclear trafficking, a step that is required for efficient gRNA packaging [7,8]. Mutants of Gag with reduced nuclear trafficking have lower levels of gRNA incorporation, whereas enhancing nuclear localization restores gRNA packaging [8]. After particle release, RSV Gag is cleaved into MA, p2, p10, CA, NC, and small spacer peptides. The mature NC protein plays important roles during the early stages of infection.

Nuclear trafficking of RSV Gag is mediated by nuclear localization signals (NLSs) in the MA and NC domains [9]. The NC region contains a classical nuclear localization signal (NLS) consisting of basic residues that bind directly to the major cellular nuclear import factor importin-alpha, which then recruits importin-beta for nuclear translocation of Gag [9,10]. In the context of NC alone, this NLS also acts as a nucleolar localization signal, with the majority of NC localizing within nucleoli [11]. The NLS in the Gag MA domain also contributes to nuclear transport, although it interacts with two different host importins [9]. A nuclear export signal (NES) in the Gag p10 domain functions through its interaction with CRM1/RanGTP export complex. Mutations of the p10 domain or treatment with leptomycin B, a CRM1 inhibitor, result in accumulation of Gag in the nucleus with formation of numerous nucleoplasmic and nucleolar foci [11–14]. These foci are dependent on the presence of the NC domain and its nucleic-acid binding function [11,14,15].

The RSV Gag NC domain contains two Cys-His domains, or zinc knuckles, that bind RNA and are required for specific gRNA packaging [16–19]. The basic residues in the Gag NC domain play numerous roles in virus assembly, including promoting Gag-Gag interactions leading to dimer and oligomer formation, nonspecific and specific nucleic acid binding, and gRNA encapsidation [20–23]. The RSV NC domain contains sixteen basic residues, however only eight are required to promote Gag-Gag interactions and virus particle assembly [21]. Lee *et al*. performed several studies examining basic residues in NC [21,22]. When the RKR residues immediately following the second zinc knuckle were deleted, they found little evidence for binding of NC to the MΨ RNA using a yeast three-hybrid assay [22]. In their follow up work, they observed that the RKR deletion mutant was still able to undergo Gag-Gag interactions [21], suggesting importance of these residues in MΨ binding, but not in Gag-Gag interactions.

To further examine roles the C-terminal basic residues in NC play in Gag subcellular localization, virus budding, gRNA packaging, and infectivity, we examined deletions and substitutions of basic residues derived from heterologous viral proteins with roles similar to NC. For this purpose, we chose the highly basic region (BR) of the herpes simplex type 1 (HSV-1) ICP27 protein, which contains an RGG Box RNA binding domain [24]. ICP27 localizes to nuclear foci and nucleoli [25–27] interacts with splicing components in nuclear speckles, [28] and binds intronless HSV-1 RNAs for nuclear export [26]. As a second viral RNA binding protein, we chose the HIV-1 Rev protein, which also localizes to nucleoli, contains an RNA binding domain enriched in basic residues, and binds to the Rev-response element to facilitate export of unspliced viral RNA from the nucleus [29–31].

## 2. Materials and Methods

### 2.1. Expression vectors, plasmids, and cells

The Prague C RSV Gag expression vector containing YFP (pGag-YFP) fluorophore was previously described [14]. RSV NC was expressed from a pEYFP-N1 containing vector (Clontech) described previously [9]. Proviral constructs were created by site-directed mutagenesis in the NC domain of pCMV.GagPol (kind gift of Rebecca Craven, Penn State College of Medicine). To create the pRS.V8.Gag.Δ61-73, pRS.V8.Gag.ICP27, and pRS.V8.Gag.Rev proviral constructs, the NC region of each Gag mutant in pCMV.GagPol was inserted into pRS.V8 using *SbfI-HpaI* restriction sites. The pGag.Δ61-73-YFP construct was made using Q5 site directed mutagenesis (New England Biolabs). To make the pGag.ICP27-YFP and pGag.Rev-YFP constructs, the NC region from pCMV.GagPol was exchanged with the NC coding sequence from pGag-YFP. Endonuclease digestion was used to identify clones containing the mutations and all positive clones were confirmed using DNA sequence analysis. All experiments were performed using the quail fibroblast QT6 cell line or the chicken fibroblast DF1 cell line [32,33]. Transfections were performed using the calcium phosphate method [34].

### 2.2. Immunofluorescence

QT6 cells seeded onto a 1.5 mm glass coverslip were transfected with wildtype or mutant provirus plasmids overnight, culture media was removed, and cells were fixed using 2% paraformaldehyde (PFA) in phosphate buffered saline (PBS) supplemented with 5 mM EGTA and 4 mM MgCl_2_, and adjusted to pH 7.2-7.4 with HCl or 3.7% PFA in 2× PHEM buffer (3.6% PIPES, 1.3% HEPES, 0.76%EGTA, 0.198% MgSO4, pH to 7.0 with 10M KOH) [35]. Cells were permeablilized using 100% methanol at RT for 5 minutes, and subsequently blocked with 5% goat serum (Rockland). After one hour, the cells were washed using 0.1% Tween-20 in PBS, and incubated with a rabbit *α*-RSV antibody (1:300) [36] and a Cy3-conjugated α-rabbit secondary antibody (1:100, Abcam). DAPI was added at 5 μg/ml. Coverslips were mounted on slides using SlowFade reagent (Invitrogen) and imaged using a Leica AOBS SP2 confocal microscope with Cy3 excited at 543nm and DAPI excited at 405nm.

### 2.3. Confocal imaging

For pNC.YFP constructs co-expressed with pFibrillarin.CFP, 0.2 × 10^6^ cells were seeded onto 35-mm glass-bottomed dishes (MatTek Corporation) and imaged using confocal microscopy at 14 to 24 h post-transfection. Sequential scanning settings were used to differentiate CFP (excitation at 458 nm, emission at 465-490 nm, and 50% laser power) and YFP (excitation at 514 nm, emission at 530-600 nm, and 10% laser power) emission spectra. The Gag-YFP wildtype and mutant proteins were imaged on the Leica SP8 confocal with DAPI excited with the 405 nm UV laser at 20% laser power using a PMT detector and YFP imaged using the WLL with a laser line excitation of 514 nm using a hybrid detector.

### 2.4. Budding analysis

Budding assays were performed as previously described in detail in [37]. Briefly, Gag expression within cell lysates was labelled for 5 minutes using ^35^S-Met/Cys. Lysates were collected then Gag was immunoprecipitated using an *α*-RSV antibody and were resolved by sodium dodecyl sulfate-polyacrylamide gel electrophoresis (SDS-PAGE) and quantified using a PhosphorImager (Bio-Rad). After ^35^S-Met/Cys labeling for 2.5-hours, supernatants were clarified, and virus particles were pelleted and immunoprecipitated using an *α*-RSV antibody and separated by SDS-PAGE and imaged by phosphorimaging. Budding efficiency was calculated as a ratio of the CA present in the media divided by the total amount of Gag expressed in the lysates. The release of the wildtype RS.V8 was set at 100%, and all mutants were expressed as a percentage of the wildtype level of budding

### 2.5. Ribonuclease protection assays

QT6 cells were transfected with either wildtype or mutant proviral DNA constructs. Culture media was collected after 48 hours, cells were pelleted using a low speed spin, and the media was passed through a 0.2 μm filter. Virus particles were pelleted by ultracentrifugation at 126,000 X g through a 25% sucrose cushion. After resuspension of the pellet, aliquots were removed for reverse transcriptase (RT) assays. The mean RT values were used to normalize the amount of virus particles for each sample as previously described [38]. Viral RNA was extracted using a QiaAMP viral RNA mini kit (Qiagen). Ribonuclease protection assays were performed as previously described [38]. A 318-nucleotide antisense probe transcribed with T7 RNA polymerase with [^32^P]CTP, spanning the splice acceptor site of the *env* gene, was used to detect both unspliced (263-nucleotide fragment) and spliced (183-nucleotide fragment) viral RNA as previously described in detail [39]. The unprotected fragments of the RNAs were digest with RNase and the samples were separated by gel electrophoresis and quantified using a PhosphorImager (BioRad).

### 2.6. Viral infectivity assay

Infection assays were performed as previously described in detail [37]. Supernatants were collected from QT6 cells expressing either wildtype or mutant provirus after 48 hours. Virus particles were concentrated by ultracentrifugation at 126,000 X *g* through a 25% sucrose cushion. RT assays were performed, as described above, to normalize the amount of virus used for infection. Equivalent amounts of RT counts from each concentrated virus preparation were added to naive DF1 cells. Cells were then assayed for the presence of GFP, which is expressed from the RS.V8 provirus, by flow cytometry (FACSCanto, BD Biosciences). The percent of cells expressing GFP was measured every three days until all infectious virus constructs reached approximately 95% green cells, or after 21 days for all remaining viruses. A minimum of three infectivity assays was performed for each proviral construct from two separate transfections.

## 3. Results

### 3.1. Substitution of the ICP27 or Rev BR for basic residues in NC altered the budding of virus particles

To examine whether the basic residues after the second Cys-His box in RSV Gag NC were important for virus assembly, we deleted residues 61-73 in the context of the proviral construct pRS.V8 (pRS.V8.Δ61-73) (Figure 1A), and a quantitative radioimmunoprecipitation assay was performed. A representative experiment showing Gag expression within the cell lysates after a 5-minute labeling period is presented in Figure 1B. After detection of metabolically-labeled Gag proteins released into the supernatant, quantitation of virus release was performed by dividing the amount of CA in the media by the amount of Gag in the cell lysates [37]. For RS.V8.Δ61-73, we observed that budding was not significantly changed compared to wild type Gag (Figure 1C and D). To determine whether budding would be affected by replacing the basic residue-containing region between amino acids 61-73 with a stretch of basic amino acids from the viral RNA binding proteins HIV-1 Rev and HSV-1 ICP27, these heterologous BRs were inserted into this region of NC [40,41], forming the chimeric proviral constructs pRS.V8.Rev and pRS.V8.ICP27 (Figure 1A).

**Figure 1.**
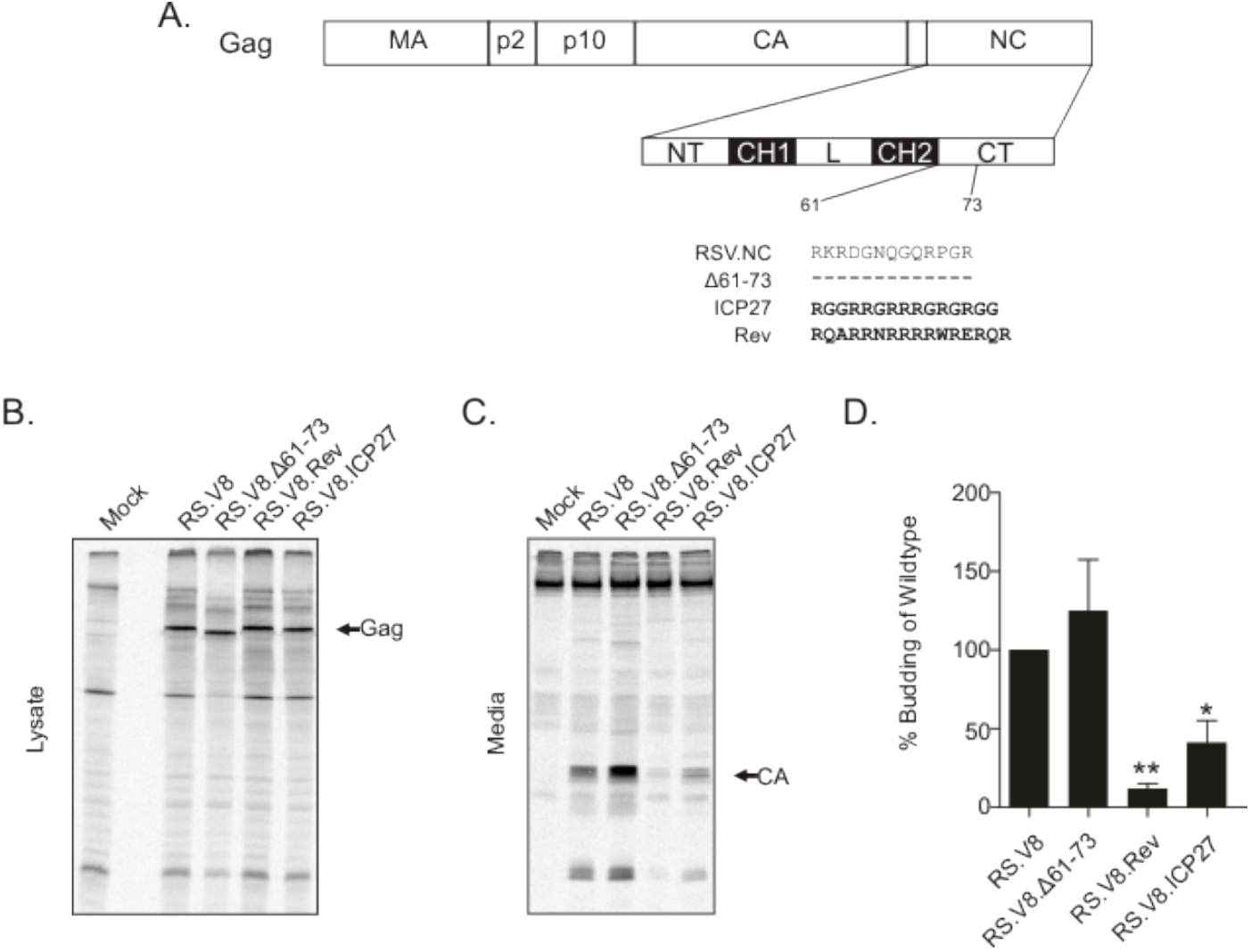
Budding analysis of proviral BR restoration mutants. (A) Schematic diagram of RSV NC and nucleolar restoration mutants. MA, matrix; CA, capsid; NC, nucleocapsid; NT, N-terminal region; CH1, Cys-His box 1; L, linker region; CH2, Cys-His box 2; CT, C-terminal region. Wildtype residues are shown for the C-terminal basic region. Dashed lines represent deleted residues (Δ61-73). Bold residues depict sequences used to replace the deleted amino acids (ICP27 BR and Rev BR). (B) Gag expression within the cell lysates after a 5-minute labeling period with ^35^S-Met/Cys. After collection of cell lysates and immunoprecipitation using an *α*-RSV antibody, the proteins were separated by SDS-PAGE and visualized by phosphorimaging. Differences in molecular weights (predicted masses listed in parentheses) were noticeable, with Δ61-73 mutant Gag (73 kD) running faster than wildtype (74.5 kD), and both Rev (75.3 kD) and ICP27 (74.7 kD) running slightly slower in the gel. The arrow indicates the position of the Gag band. (C) After radioactive labeling for 2.5-hours, supernatants were clarified, and virus particles were pelleted and immunoprecipitated using an *α*-RSV antibody (media samples). Proteins were separated by SDS-PAGE and visualized by phosphorimaging. The arrow indicates CA (25.8 kD). (D) The average of four independent budding assays is presented within the bar graph, with the bars representing the standard error. *p value = 0.0253, **p value = 0.0013 calculated by Student’s t-test.

To determine whether insertion of the BRs from ICP27 and Rev altered virus particle production, budding efficiency for each mutant was compared to wildtype, which was set at 100% (Figure 1D). RS.V8.Δ61-73 was not significantly different from wildtype (147%, p > 0.5). RS.V8.Rev was reduced in budding (12%; p = 0.0013), as was RS.V8.ICP27 (41%; p = 0.0253). These results indicate that the BR from Rev and ICP27 negatively affects budding, but the complete removal of the BR in the C-terminal region of NC (RS.V8.Δ61-73) does not.

### 3.2. Subcellular localization of the ICP27 BR and Rev BR Gag mutants

The amino acid sequences derived from ICP27 and Rev are capable of general RNA binding as well as facilitating nucleolar localization [24–27,29–31]. Therefore, we tested whether the addition of these heterologous viral RNA binding domains, which also serve as nucleolar localization signals, would alter the normal trafficking patterns of RSV Gag. To investigate this possibility, we examined cells expressing wildtype or mutant proviral constructs by immunofluorescence staining using a polyclonal *α*-RSV antibody (Figure 2A). The wildtype Gag protein expressed using a proviral vector exhibited a small amount of nuclear fluorescence, with the majority of the fluorescence signal in the cytoplasm, forming discrete foci at the plasma membrane (Figure 2A, panel a). The Δ61-73 mutant showed similar localization, with fluorescent signal primarily in the cytoplasm and along the plasma membrane and possibly even more strongly at the plasma membrane compared to wildtype (Figure 2A, panel b). The Gag localization results correlate with the budding data from Figure 1, in that the average budding efficiency of Δ61-73 was greater than wildtype, even though not statistically significant. Unexpectedly, although both RS.V8 ICP27 and Rev mutants contain the NES in the Gag p10 domain, they both were concentrated in the nucleus and formed numerous foci in the nucleoplasm (Figure 2A, panels c and d, respectively).

**Figure 2.**
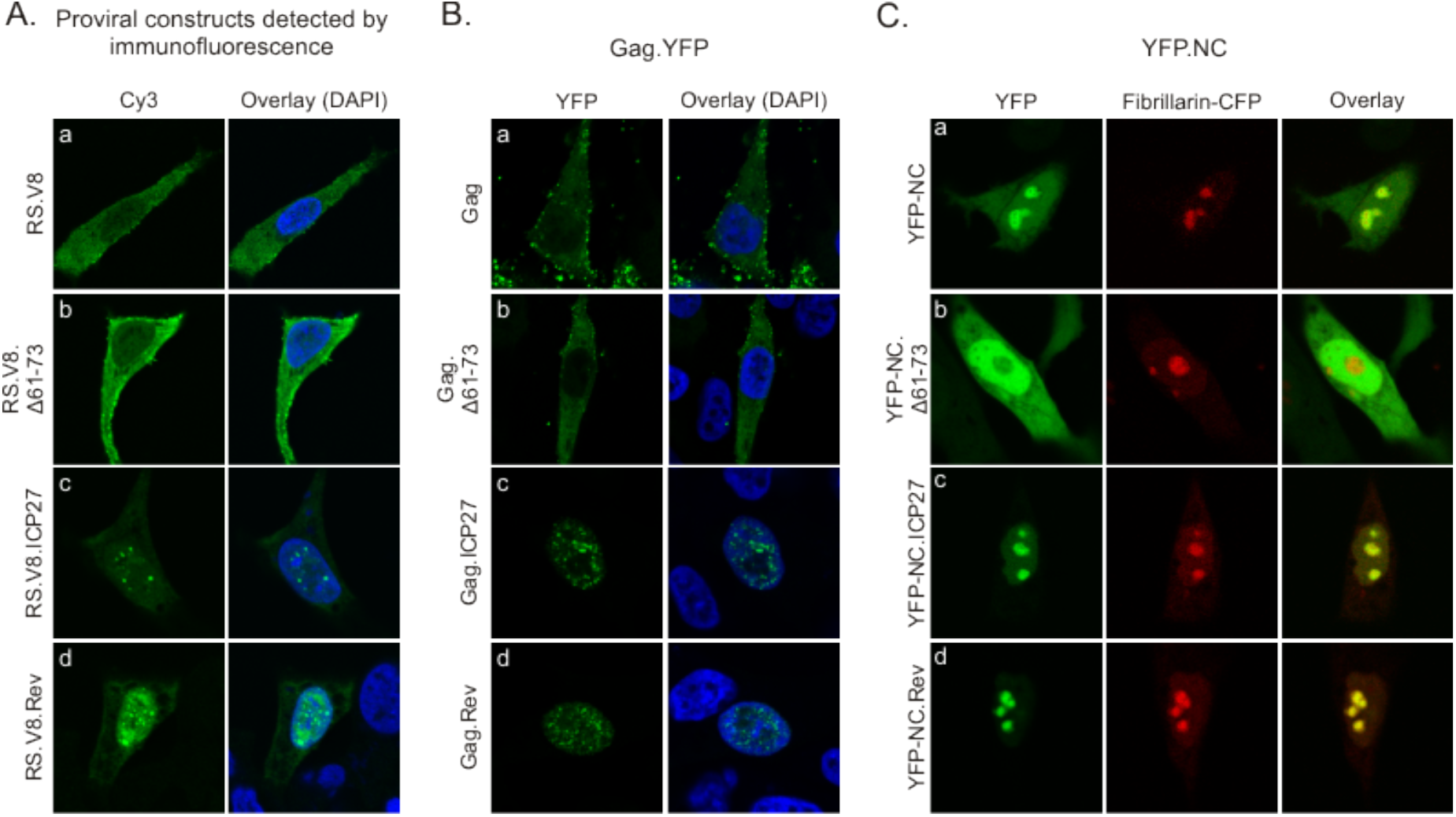
Subcellular localization of Gag ICP27 BR and Rev BR substitution mutants. (A) QT6 cells expressing wildtype or mutant proviral constructs were immunostained with anti-RSV antibodies 16 hours after transfection to detect localization of Gag. Confocal microscopy images taken through the central plane of the nucleus show the distribution of wildtype Gag (panel a), Gag.Δ61-73 (panel b), Gag.ICP27 (panel c) and Gag.Rev (panel d). (B) Plasmids encoding wildtype (panel a) and mutant Gag.YFP proteins (panels b-d) were expressed in QT6 cells and fluorescence was detected using confocal microscopy. (C) QT6 cells were co-transfected with wildtype YFP-NC (panel a; green) or mutant YFP-NC proteins (panels b-d; green) and fibrillarin-CFP (red) as a marker for nucleoli, with the overlay images showing colocalization (yellow).

To determine whether viral nucleic acids or proteins expressed from the proviral vectors affected Gag localization, we expressed Gag fused to YFP using a CMV promoter (Figure 2B). Gag.YFP was similar in appearance, with mostly cytoplasmic and plasma membrane fluorescence and a lower amount of diffuse and focal nuclear signals (Figure 2B, panel a). The deletion mutant Gag.Δ61-73 had similar localization compared to wildtype Gag (Figure 2B, panel b). However, when either the Gag.ICP27.YFP or Gag.Rev.YFP mutants were expressed, they localized almost exclusively to the nucleus, forming numerous nucleoplasmic foci (Figure 2B, panels c and d, respectively). Of note, full length ICP27 forms nucleoplasmic foci in HSV-infected cells [24–26], whereas HIV-1 Rev localizes to nucleoli under steady-state conditions [30].

When expressed by itself, wildtype NC localizes primarily to nucleoli [11] (Figure 2C, panel a). By contrast, the NC.Δ61-73 deletion mutant was nuclear-localized but excluded nucleoli (Figure 2C, panel b). Substitution of the ICP27 or Rev BRs for residues 61-73 restored nucleolar localization of the NC protein (Figure 2C, panels c and d, respectively). These data indicate that replacement of amino acids 61-73 with either the ICP27 or Rev BR is sufficient to direct NC nucleolar localization.

### 3.3. Heterologous BRs substituted in the Gag NC domain restore gRNA packaging

The Cys-His boxes in RSV Gag specifically bind to the psi-packaging sequence [17,18,42], although basic residues within NC also contribute to gRNA packaging [21,22]. To investigate the importance of basic residues in the C-terminal region of the Gag for gRNA encapsidation, we measured the relative amount of gRNA in virus particles using a quantitative ribonuclease protection assay. The amount of gRNA isolated from wildtype and mutant viruses was normalized using reverse transcriptase activity, as described in Materials and Methods. We used a probe spanning the 3’ splice acceptor site in *env* to quantitate the amount of spliced viral RNA and unspliced gRNA isolated from purified particles. Representative autoradiograms are shown (Figure 3A), and the means of at least three independent experiments for each mutant were plotted (Figure 3B). The amount of gRNA detected in wildtype virions was set to 100%, and each mutant was compared to wildtype. The Gag.Δ61-73 mutant virus was significantly reduced in its ability to package gRNA (15% of the wildtype level, p = 0.0001). The RS.V8.Rev and RS.V8.ICP27 viruses packaged RSV gRNA much more efficiently, with levels of 73% and 78% compared to wildtype, respectively, although gRNA incorporation for RS.V8.Rev was statistically lower than wildtype (p = 0.0278). These results indicate that insertion of a heterologous BR restores incorporation of gRNA into virus particles at near wildtype levels. When the ratio of spliced:unspliced viral RNA were examined, both BR insertion mutants packaged nearly the same amount as wildtype virus. However, in the case of the Δ61-73 mutant virus, the level of packaging of gRNA was drastically reduced, and the ratio of spliced:unspliced viral RNA was increased compared to wildtype to nearly 1:1, demonstrating a reduction in the specificity of the type of viral RNA incorporated into particles.

**Figure 3.**
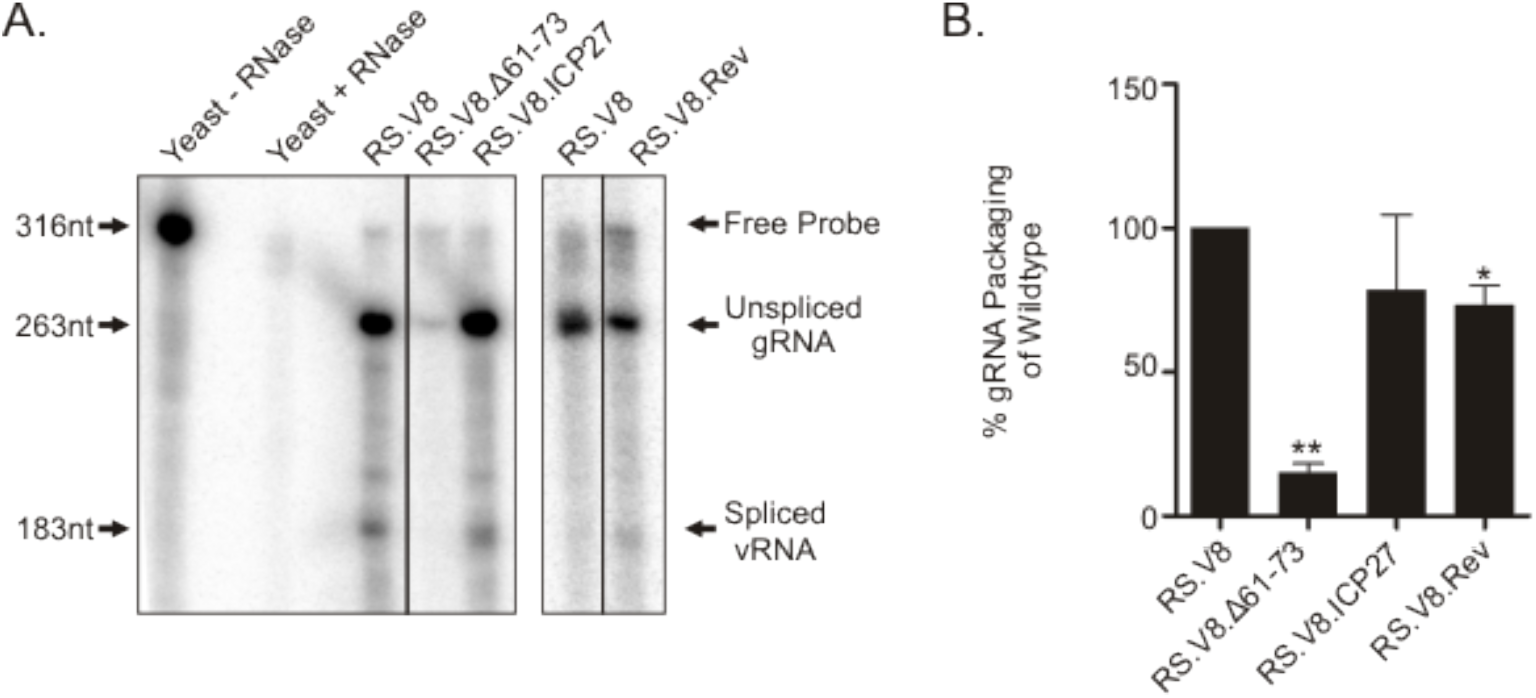
gRNA packaging efficiency of heterologous BR mutants. (A) Wildtype RS.V8 or mutant proviral vectors containing the Gag.Δ61.73 deletion or substitutions of Gag.ICP27 BR or Gag.Rev BR were transfected into QT6 cells. Virus particles were collected for 48 hours and were normalized using reverse transcriptase assay. Equivalent amounts of virus particles were used to extract RNA, and ribonuclease protection assay was used to measure the levels of spliced and unspliced viral RNA present. After hybridization of the 318-nt ^32^P-radiolabled antisense riboprobe, which spans the 3’-splice acceptor site in *env*, and digestion of the unprotected fragment with RNase treatment, the RNA was separated by gel electrophoresis and visualized using a phosphorimager. Black lines represent lanes that were removed and spaces between the gels represent results from independent experiments. Arrows to the left identify nucleotide lengths, and arrows to the right denote the species of RNA (free probe, unspliced gRNA, and spliced viral RNA). (B) The results of four independent experiments are shown on the graph, with the bars representing standard error of the mean. * p value = 0.0278; ** p value = 0.0001 when compared to wildtype levels by Student’s t-test.

### 3.4. Viruses with deletions of basic residues or heterologous BR substitutions in the NC domain of Gag are noninfectious

To determine whether the basic residues in the C-terminal region of Gag are required for virus replication, we performed infectivity assays. Virus particles were collected from QT6 cells expressing either wildtype or mutant proviral constructs, the particles were normalized by reverse transcriptase activity, and equivalent amounts of virus were placed on uninfected DF1 cells. The RS.V8 provirus contains a GFP gene in the place of *src*, so the ability of mutant viruses to spread through the cell culture was monitored by measuring fluorescence using fluorescence activated cell every three days for 21 days (Figure 4).

**Figure 4.**
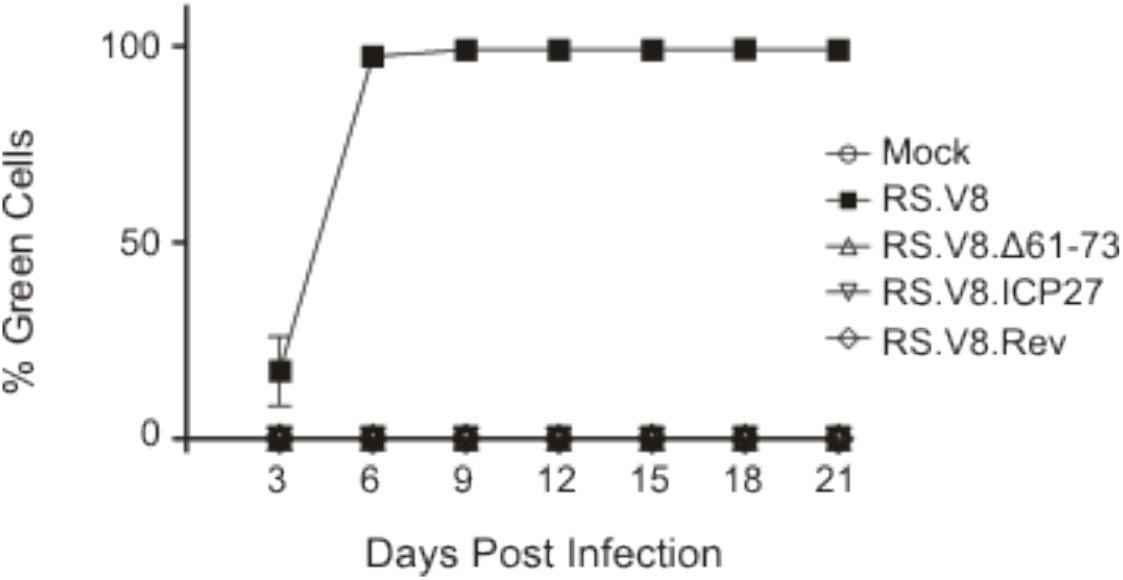
Analysis of infectivity. Virus particles were collected from QT6 cells transfected with wildtype pRS.V8 or mutant proviruses, as indicated. Particles were normalized and then used to infect naive DF1 cells. Infected cells were passaged and examined every 3 days post-infection to detect expression of GFP from integrated proviruses using flow cytometry. The percentage of GFP-expressing cells at each time point was plotted. Neither the deletion mutant (Δ61-73) nor either of the substitution mutant viruses were infectious. The graph shown is the average of three independent infection assays with error bars showing standard error of the mean.

The wildtype virus (RS.V8) was able to efficiently infect and replicate within cells, reaching approximately 95% of cells expressing the GFP protein by day 6 post-infection. The RS.V8.Δ61-73 mutant was noninfectious, which was expected given the low incorporation of gRNA. However, even though gRNA incorporation was restored by insertion of the exogenous viral RNA binding domains from ICP27 and Rev, these substitutions were unable to restore infectivity. We cannot distinguish between the possibilities that these mutant viruses are defective in establishment of infection and/or cell-to-cell spread.

## 4. Discussion

The NC domain of the RSV Gag protein is enriched in basic residues, which play a variety of roles in virus replication [20–22]. To date, the functions of basic residues in NC have been primarily studied using deletion mutagenesis, which produces negative or loss-of-function results. More informative are genetic gain-of-function experiments, in which an essential function is reconstituted using functional domains derived from heterologous proteins. In this report, we first deleted a region following the Cys-His boxes of RSV NC that contains five positively charged residues, finding that Gag localization and budding were not adversely affected, whereas gRNA packaging and infectivity were severely compromised, and nucleolar localization of the mature NC protein was abrogated. The data suggest that these basic residues do not affect general Gag-nucleic acid interactions which are needed for particle assembly [43–45], but appear to be involved in the specificity of binding viral gRNA.

In an attempt to separate the contributions of basic residues encompassing amino acids 61-73 in RSV NC, we substituted the sequences from two different viral RNA binding proteins [27,46] that also contain nucleolar localization signals. Even though these sequences were derived from very different viral proteins, those of HSV-1 ICP27 and HIV-1 Rev, the resulting chimeric proteins behaved very similarly. The most striking feature was the re-localization of these chimeric Gag proteins to the nucleus, with accumulation in discrete nucleoplasmic foci that resembled ICP27 nuclear foci in HSV-1 infected cells [28,47,48]. In fact, quite surprisingly, the ICP27 and Rev Gag nuclear foci are very similar in appearance to RSV Gag treated with leptomycin B (an inhibitor of Crm1 nuclear export) or with the introduction of a point mutant in the p10 NES of Gag that blocks nuclear egress [14,15].

What accounts for the accumulation of Gag in nuclear foci with the addition of BRs from Rev and ICP27? The NC domain of Gag is a major site of Gag-Gag interactions mediated through protein-protein or protein-RNA binding [23,49,50], so perhaps the BRs enhance Gag-Gag interactions. We previously showed that Gag-Gag intermolecular interactions in the nucleus depend on the NC domain and its RNA binding ability [14]. In addition, the ICP27 and Rev RNA binding domains may interact with host nuclear factors, either proteins and/or RNAs, that serve to tether Gag more strongly in the nucleus [28,47,51–54]. We previously reported that RSV Gag colocalizes with splicing factors [15], as does ICP27[28]. Whether Gag influences RNA splicing, as has been found for ICP27 [51], is not yet known. It is also possible that the insertions of ICP27 and Rev BRs in NC alter the conformation of the Gag protein, interfering with the function of the NES in p10, leading to nuclear retention of the mutant Gag proteins.

The increased nuclear localization of the chimeric Gag proteins likely explains the budding defect of the RS.V8.ICP27 and RS.V8.Rev viruses. These results provide further evidence that nuclear trafficking of RSV Gag is intrinsic to the virus assembly pathway since enhancing nuclear localization was linked to an assembly defect these mutants. In spite of the defect in particle production, the ICP27 and Rev chimeric viruses both restored gRNA packaging when compared to mutant bearing a deletion of basic residues within 61-73 of the NC domain. This restoration of gRNA packaging could be due to the RNA binding capabilities of the RS.V8.Rev and RS.V8.ICP27 mutants, although it was somewhat unexpected that the RNA binding domains from heterologous viruses would confer selective packaging of the RSV genome, maintaining the proper ratio of spliced:unspliced viral RNAs incorporated into virus particles.

One possible explanation for the effect of the BR insertions is that the presence of several nonspecific RNA-binding domains may contribute to specific gRNA binding, although the mechanisms underlying specificity of RNA recognition is not fully understood [55]. For example, RNA binding domains rich in RS or RG residues, such as the RGG box of ICP27, in combination with other basic sequences, have been shown to mediate both specific and non-specific interactions with RNA [56]. The properties of RNA binding proteins have been extensively studied in serine and arginine-rich (SR) proteins involved in RNA splicing [56,57]. SR proteins generally contain one or two of the RNA recognition motifs at the N-terminus and a RS motif at the C-terminus. Typically, the RNA recognition motifs are involved in recognizing the specific target RNAs, while the RS motifs participate in indirect protein-protein and non-specific protein-RNA interactions that bring the SR proteins and the target RNAs within close proximity [56,57]. Other common sequences typically found in RNA binding proteins are motifs enriched in R/K residues, in which four to eight residues in small patches form highly positive regions that mediate molecular interactions. They frequently flank globular domains, assisting in RNA binding [56]. Therefore, it is possible that the deletion of basic residues immediately after the second Cys-His box in RSV Gag eliminates specific binding of the Ψ-sequence, as previously suggested by Lee et al. [22]. Introduction of additional basic residues in the ICP27 and Rev heterologous RNA-binding domains, which contain 8 and 10 basic residues, respectively, restores the interaction with RSV RNA via electrostatic interactions that bring Gag and gRNA in close proximity, allowing specific gRNA binding through the zinc knuckles in NC.

Electrostatic interactions play a role in the binding of positively charged RNA binding proteins to negatively charged RNAs through both specific and non-specific mechanisms. We searched the literature for cellular RNA binding proteins with mechanisms that explain how non-specific electrostatic interactions could contribute to specific binding to their cognate RNA molecules. We found the spliceosomal protein, U1A uses a non-specific ‘lure’ step followed by a specific ‘lock’ step to binds to its SL2 RNA partner (summarized in [58]). Short-range electrostatic interactions during the ‘lure’ step, are proposed to attract the SL2 RNA to U1A, followed by the ‘lock’ step, in which long-range electrostatic interactions between the protein and RNA create specific binding. We hypothesize that RSV Gag could be utilizing a similar technique for binding to viral gRNA. The basic residues in the C-terminus of NC could be involved in the initial electrostatic ‘lure’ step, bringing gRNA in closer proximity. This binding event would then allow the zinc knuckles in NC to specifically bind to the psi sequence on the viral RNA. Thus, whether selective gRNA packaging in the chimeric ICP27 and Rev viruses is due to the addition of these heterologous RNA binding domains or simply the addition of additional basic amino acids is not clear. Further studies to examine this question would be needed, perhaps by inserting a random sequence of basic residue sequences versus RNA binding domains that do not contain basic amino acids (KH domain or additional zinc fingers) to further dissect the mechanism.

Although insertion of heterologous BRs from ICP27 and Rev into NC rescued selective incorporation of gRNA into virus particles, these mutants were noninfectious. Because the mutations were inserted into NC, the most likely explanation is that functions of NC in other steps of infection were impaired, including gRNA dimerization [16,59,60], reverse transcription [61–65], transport of the preintegration complex [66], or chaperone function [67,68]. We can conclude that nucleolar localization of NC, which was restored in the ICP27 and Rev chimeras, is not sufficient for NC-mediated replication activities early in infection. Further investigation into the mechanisms underlying the replication defect of these chimeric viruses will be enlightening.

## 5. Conclusions

The work presented here demonstrates the importance of the basic residues after the Cys-His boxes in the C-terminus of RSV NC in gRNA packaging. When this sequence is deleted, virus particles are able to bud normally from cells, but are not infectious. However, when these residues are replaced with heterologous basic residues from other viral RNA binding proteins, the subcellular localization of Gag became predominantly nuclear, and gRNA packaging was restored. Even though gRNA packaging in the chimeras was increased to near normal levels, these viruses remained noninfectious. These results suggest that the C-terminal basic residues in NC are important for facilitating gRNA binding. A deeper understanding of the mechanism by which retroviral Gag proteins selectively incorporate their genomes may be helpful in future antiviral and vaccine development [69–73].

## Author Contributions

B.L.R. Writing – original draft preparation, conceptualization, formal analysis, investigation; T.L.L. Conceptualization, formal analysis, investigation, methodology, writing – original draft and review and editing; L.J.P. Conceptualization, funding acquisition, project supervision, resources, writing— review and editing. All authors have read and agreed to the published version of the manuscript.

## Funding

This research was funded by the National Institute of Health grants: R01 CA76534 (LJP), T32 CA60395 (TLL), F31 CA196292 (BLR), and internal funding from the Department of Medicine and the Penn State College of Medicine.

## Acknowledgments

We would like to thank Dr. Rebecca Craven (Penn State College of Medicine) for generosity in supplying reagents. We would like to acknowledge the Microscopy Imaging Core Facility at PSU College of Medicine for use of the Leica AOBS SP2 confocal and the Leica SP8 confocal [1S10OD010756-01A1 (CB)].

## Conflicts of Interest

The authors declare no conflict of interest.

## References

1. Kaddis Maldonado, R.J.; Parent, L.J. Orchestrating the Selection and Packaging of Genomic RNA by Retroviruses: An Ensemble of Viral and Host Factors. Viruses 2016, 8.

2. Muriaux, D.; Darlix, J.L. Properties and functions of the nucleocapsid protein in virus assembly. RNA Biol 2010, 7, 744–753, doi:10.4161/rna.7.6.14065.

3. Darlix, J.L.; de Rocquigny, H.; Mauffret, O.; Mely, Y. Retrospective on the all-in-one retroviral nucleocapsid protein. Virus Res 2014, 193, 2–15, doi:10.1016/j.virusres.2014.05.011.

4. Mirambeau, G.; Lyonnais, S.; Gorelick, R.J. Features, processing states, and heterologous protein interactions in the modulation of the retroviral nucleocapsid protein function. RNA Biol 2010, 7, 724–734, doi:10.4161/rna.7.6.13777.

5. Olson, E.D.; Musier-Forsyth, K. Retroviral Gag protein-RNA interactions: Implications for specific genomic RNA packaging and virion assembly. Semin Cell Dev Biol 2019, 86, 129–139, doi:10.1016/j.semcdb.2018.03.015.

6. Zhou, J.; McAllen, J.K.; Tailor, Y.; Summers, M.F. High affinity nucleocapsid protein binding to the muPsi RNA packaging signal of Rous sarcoma virus. J Mol Biol 2005, 349, 976–988, doi:10.1016/j.jmb.2005.04.046.

7. Scheifele, L.Z.; Rhoads, J.D.; Parent, L.J. Specificity of plasma membrane targeting by the rous sarcoma virus gag protein. J.Virol. 2003, 77, 470–480.

8. Garbitt-Hirst, R.; Kenney, S.P.; Parent, L.J. Genetic evidence for a connection between Rous sarcoma virus gag nuclear trafficking and genomic RNA packaging. J.Virol. 2009, 83, 6790–6797.

9. Butterfield-Gerson, K.L.; Scheifele, L.Z.; Ryan, E.P.; Hopper, A.K.; Parent, L.J. Importin-beta family members mediate alpharetrovirus gag nuclear entry via interactions with matrix and nucleocapsid. J.Virol. 2006, 80, 1798–1806.

10. Gudleski, N.; Flanagan, J.M.; Ryan, E.P.; Bewley, M.C.; Parent, L.J. Directionality of nucleocytoplasmic transport of the retroviral gag protein depends on sequential binding of karyopherins and viral RNA. Proc Natl Acad Sci U S A 2010, 107, 9358–9363, doi:10.1073/pnas.1000304107.

11. Lochmann, T.L.; Bann, D.V.; Ryan, E.P.; Beyer, A.R.; Mao, A.; Cochrane, A.; Parent, L.J. NC-mediated nucleolar localization of retroviral gag proteins. Virus research 2013, 171, 304–318.

12. Scheifele, L.Z.; Garbitt, R.A.; Rhoads, J.D.; Parent, L.J. Nuclear entry and CRM1-dependent nuclear export of the Rous sarcoma virus Gag polyprotein. Proc.Natl.Acad.Sci.U.S.A 2002, 99, 3944–3949.

13. Scheifele, L.Z.; Ryan, E.P.; Parent, L.J. Detailed mapping of the nuclear export signal in the Rous sarcoma virus Gag protein. J.Virol. 2005, 79, 8732–8741.

14. Kenney, S.P.; Lochmann, T.L.; Schmid, C.L.; Parent, L.J. Intermolecular interactions between retroviral Gag proteins in the nucleus. J.Virol. 2008, 82, 683–691.

15. Rice, B.L.; Kaddis, R.J.; Stake, M.S.; Lochmann, T.L.; Parent, L.J. Interplay between the alpharetroviral Gag protein and SR proteins SF2 and SC35 in the nucleus. Front Microbiol 2015, 6.

16. Bowles, N.E.; Damay, P.; Spahr, P.F. Effect of rearrangements and duplications of the Cys-His motifs of Rous sarcoma virus nucleocapsid protein. J.Virol. 1993, 67, 623–631.

17. Meric, C.; Gouilloud, E.; Spahr, P.F. Mutations in Rous sarcoma virus nucleocapsid protein p12 (NC): deletions of Cys-His boxes. J.Virol. 1988, 62, 3328–3333.

18. Zhou, J.; Bean, R.L.; Vogt, V.M.; Summers, M. Solution structure of the Rous sarcoma virus nucleocapsid protein: muPsi RNA packaging signal complex. J.Mol.Biol. 2007, 365, 453–467.

19. Dupraz, P.; Oertle, S.; Meric, C.; Damay, P.; Spahr, P.F. Point mutations in the proximal Cys-His box of Rous sarcoma virus nucleocapsid protein. J.Virol. 1990, 64, 4978–4987.

20. René, B.; Mauffret, O.; Fossé, P. Retroviral nucleocapsid proteins and DNA strand transfers. Biochimie open 2018, 7, 10–25, doi:10.1016/j.biopen.2018.07.001.

21. Lee, E.G.; Linial, M.L. Basic residues of the retroviral nucleocapsid play different roles in gag-gag and Gag-Psi RNA interactions. J.Virol. 2004, 78, 8486–8495.

22. Lee, E.G.; Alidina, A.; May, C.; Linial, M.L. Importance of basic residues in binding of rous sarcoma virus nucleocapsid to the RNA packaging signal. J.Virol. 2003, 77, 2010–2020.

23. Bowzard, J.B.; Bennett, R.P.; Krishna, N.K.; Ernst, S.M.; Rein, A.; Wills, J.W. Importance of basic residues in the nucleocapsid sequence for retrovirus Gag assembly and complementation rescue. J Virol 1998, 72, 9034–9044.

24. Lengyel, J.; Guy, C.; Leong, V.; Borge, S.; Rice, S.A. Mapping of functional regions in the amino-terminal portion of the herpes simplex virus ICP27 regulatory protein: importance of the leucine-rich nuclear export signal and RGG Box RNA-binding domain. J Virol 2002, 76, 11866–11879, doi:10.1128/jvi.76.23.11866-11879.2002.

25. Mears, W.E.; Lam, V.; Rice, S.A. Identification of nuclear and nucleolar localization signals in the herpes simplex virus regulatory protein ICP27. J Virol 1995, 69, 935–947.

26. Sandri-Goldin, R.M. ICP27 mediates HSV RNA export by shuttling through a leucine-rich nuclear export signal and binding viral intronless RNAs through an RGG motif. Genes Dev 1998, 12, 868–879, doi:10.1101/gad.12.6.868.

27. Mears, W.E.; Rice, S.A. The RGG box motif of the herpes simplex virus ICP27 protein mediates an RNA-binding activity and determines in vivo methylation. J Virol 1996, 70, 7445–7453.

28. Sciabica, K.S.; Dai, Q.J.; Sandri-Goldin, R.M. ICP27 interacts with SRPK1 to mediate HSV splicing inhibition by altering SR protein phosphorylation. Embo J 2003, 22, 1608–1619, doi:10.1093/emboj/cdg166.

29. Fankhauser, C.; Izaurralde, E.; Adachi, Y.; Wingfield, P.; Laemmli, U.K. Specific complex of human immunodeficiency virus type 1 rev and nucleolar B23 proteins: dissociation by the Rev response element. Mol.Cell Biol. 1991, 11, 2567–2575.

30. Daelemans, D.; Costes, S.V.; Cho, E.H.; Erwin-Cohen, R.A.; Lockett, S.; Pavlakis, G.N. In vivo HIV-1 Rev multimerization in the nucleolus and cytoplasm identified by fluorescence resonance energy transfer. J.Biol.Chem. 2004, 279, 50167–50175.

31. Berger, J.; Aepinus, C.; Dobrovnik, M.; Fleckenstein, B.; Hauber, J.; Bohnlein, E. Mutational analysis of functional domains in the HIV-1 Rev trans-regulatory protein. Virology 1991, 183, 630–635.

32. Himly, M.; Foster, D.N.; Bottoli, I.; Iacovoni, J.S.; Vogt, P.K. The DF-1 chicken fibroblast cell line: transformation induced by diverse oncogenes and cell death resulting from infection by avian leukosis viruses. Virology 1998, 248, 295–304, doi:S0042-6822(98)99290-X [pii] 10.1006/viro.1998.9290.

33. Parent, L.J.; Cairns, T.M.; Albert, J.A.; Wilson, C.B.; Wills, J.W.; Craven, R.C. RNA dimerization defect in a Rous sarcoma virus matrix mutant. J Virol 2000, 74, 164–172.

34. Fujiwara, T.; Oda, K.; Yokota, S.; Takatsuki, A.; Ikehara, Y. Brefeldin A causes disassembly of the Golgi complex and accumulation of secretory proteins in the endoplasmic reticulum. J Biol Chem 1988, 263, 18545–18552.

35. Matic, I.; van Hagen, M.; Schimmel, J.; Macek, B.; Ogg, S.C.; Tatham, M.H.; Hay, R.T.; Lamond, A.I.; Mann, M.; Vertegaal, A.C.O. In vivo identification of human small ubiquitin-like modifier polymerization sites by high accuracy mass spectrometry and an in vitro to in vivo strategy. Mol Cell Proteomics 2008, 7, 132–144.

36. Weldon, R.A.; Erdie, C.R.; Oliver, M.G.; Wills, J.W. Incorporation of chimeric gag protein into retroviral particles. Journal of Virology 1990, 64, 4169.

37. Callahan, E.M.; Wills, J.W. Repositioning basic residues in the M domain of the Rous sarcoma virus gag protein. J.Virol. 2000, 74, 11222–11229.

38. Craven, R.C.; Leure-duPree, A.E.; Weldon, R.A. Jr.; Wills, J.W. Genetic analysis of the major homology region of the Rous sarcoma virus Gag protein. J.Virol. 1995, 69, 4213–4227.

39. Garbitt, R.A.; Bone, K.R.; Parent, L.J. Insertion of a classical nuclear import signal into the matrix domain of the Rous sarcoma virus Gag protein interferes with virus replication. J.Virol. 2004, 78, 13534–13542.

40. Mears, W.E.; Rice, S.A. The herpes simplex virus immediate-early protein ICP27 shuttles between nucleus and cytoplasm. Virology 1998, 242, 128–137, doi:S0042-6822(97)99006-1[pii] 10.1006/viro.1997.9006.

41. Bohnlein, E.; Berger, J.; Hauber, J. Functional mapping of the human immunodeficiency virus type 1 Rev RNA binding domain: new insights into the domain structure of Rev and Rex. J.Virol. 1991, 65, 7051–7055.

42. Lee, E.G.; Linial, M.L. Deletion of a Cys-His motif from the Alpharetrovirus nucleocapsid domain reveals late domain mutant-like budding defects. Virology 2006, 347, 226–233.

43. Gallis, B.; Linial, M.; Eisenman, R. An avian oncovirus mutant deficient in genomic RNA: characterization of the packaged RNA as cellular messenger RNA. Virology 1979, 94, 146–161.

44. Anderson, D.J.; Stone, J.; Lum, R.; Linial, M.L. The packaging phenotype of the SE21Q1b provirus is related to high proviral expression and not trans-acting factors. Journal of Virology 1995, 69, 7319.

45. Shank, P.R.; Linial, M. Avian oncovirus mutant (SE21Q1b) deficient in genomic RNA: characterization of a deletion in the provirus. J Virol 1980, 36, 450–456.

46. Perkins, A.; Cochrane, A.W.; Ruben, S.M.; Rosen, C.A. Structural and functional characterization of the human immunodeficiency virus rev protein. J.Acquir.Immune.Defic.Syndr. 1989, 2, 256–263.

47. Malik, P.; Tabarraei, A.; Kehlenbach, R.H.; Korfali, N.; Iwasawa, R.; Graham, S.V.; Schirmer, E.C. Herpes simplex virus ICP27 protein directly interacts with the nuclear pore complex through Nup62, inhibiting host nucleocytoplasmic transport pathways. J Biol Chem 2012, 287, 12277–12292, doi:10.1074/jbc.M111.331777.

48. Rojas, S.; Corbin-Lickfett, K.A.; Escudero-Paunetto, L.; Sandri-Goldin, R.M. ICP27 Phosphorylation Site Mutants Are Defective in Herpes Simplex Virus 1 Replication and Gene Expression. Journal of Virology 2010, 84, 2200, doi:10.1128/jvi.00917-09.

49. Wills, J.W.; Craven, R.C. Form, function, and use of retroviral gag proteins. AIDS 1991, 5, 639–654, doi:10.1097/00002030-199106000-00002.

50. Ma, Y.M.; Vogt, V.M. Rous sarcoma virus Gag protein-oligonucleotide interaction suggests a critical role for protein dimer formation in assembly. J Virol 2002, 76, 5452–5462, doi:10.1128/jvi.76.11.5452-5462.2002.

51. Sandri-Goldin, R.M. The many roles of the regulatory protein ICP27 during herpes simplex virus infection. Front Biosci 2008, 13, 5241–5256, doi:10.2741/3078.

52. Tunnicliffe, R.B.; Tian, X.; Storer, J.; Sandri-Goldin, R.M.; Golovanov, A.P. Overlapping motifs on the herpes viral proteins ICP27 and ORF57 mediate interactions with the mRNA export adaptors ALYREF and UIF. Scientific Reports 2018, 8, 15005, doi:10.1038/s41598-018-33379-x.

53. Naji, S.; Ambrus, G.; Cimermančič, P.; Reyes, J.R.; Johnson, J.R.; Filbrandt, R.; Huber, M.D.; Vesely, P.; Krogan, N.J.; Yates, J.R., 3rd, et al. Host cell interactome of HIV-1 Rev includes RNA helicases involved in multiple facets of virus production. Molecular & cellular proteomics : MCP 2012, 11, M111.015313–M015111.015313, doi:10.1074/mcp.M111.015313.

54. Modem, S.; Thipparthi, R.R. Cellular Proteins and HIV-1 Rev Function. Current HIV Research 2009, 7, 91–100, doi:http://dx.doi.org/10.2174/157016209787048474.

55. Lunde, B.M.; Moore, C.; Varani, G. RNA-binding proteins: modular design for efficient function. Nat Rev Mol Cell Biol 2007, 8, 479–490, doi:10.1038/nrm2178.

56. Balcerak, A.; Trebinska-Stryjewska, A.; Konopinski, R.; Wakula, M.; Grzybowska, E.A. RNA–protein interactions: disorder, moonlighting and junk contribute to eukaryotic complexity. Open Biology 9, 190096, doi:10.1098/rsob.190096.

57. Jeong, S. SR Proteins: Binders, Regulators, and Connectors of RNA. Mol Cells 2017, 40, 1–9.

58. Ghaemi, Z.; Guzman, I.; Gnutt, D.; Luthey-Schulten, Z.; Gruebele, M. Role of Electrostatics in Protein– RNA Binding: The Global vs the Local Energy Landscape. The Journal of Physical Chemistry B 2017, 121, 8437–8446, doi:10.1021/acs.jpcb.7b04318.

59. Baba, S.; Takahashi, K.; Koyanagi, Y.; Yamamoto, N.; Takaku, H.; Gorelick, R.J.; Kawai, G. Role of the zinc fingers of HIV-1 nucleocapsid protein in maturation of genomic RNA. J Biochem 2003, 134, 637–639.

60. Prats, A.C.; Housset, V.; de Billy, G.; Cornille, F.; Prats, H.; Roques, B.; Darlix, J.L. Viral RNA annealing activities of the nucleocapsid protein of Moloney murine leukemia virus are zinc independent. Nucleic Acids Res 1991, 19, 3533–3541.

61. Allain, B.; Lapadat-Tapolsky, M.; Berlioz, C.; Darlix, J.L. Transactivation of the minus-strand DNA transfer by nucleocapsid protein during reverse transcription of the retroviral genome. Embo J 1994, 13, 973–981.

62. Barat, C.; Schatz, O.; Le Grice, S.; Darlix, J.L. Analysis of the interactions of HIV1 replication primer tRNA(Lys,3) with nucleocapsid protein and reverse transcriptase. J Mol Biol 1993, 231, 185–190, doi:10.1006/jmbi.1993.1273.

63. Darlix, J.L.; Vincent, A.; Gabus, C.; de Rocquigny, H.; Roques, B. Trans-activation of the 5’ to 3’ viral DNA strand transfer by nucleocapsid protein during reverse transcription of HIV1 RNA. C R Acad Sci III 1993, 316, 763–771.

64. Drummond, J.E.; Mounts, P.; Gorelick, R.J.; Casas-Finet, J.R.; Bosche, W.J.; Henderson, L.E.; Waters, D.J.; Arthur, L.O. Wild-type and mutant HIV type 1 nucleocapsid proteins increase the proportion of long cDNA transcripts by viral reverse transcriptase. AIDS Res Hum Retroviruses 1997, 13, 533–543.

65. Guo, J.; Wu, T.; Anderson, J.; Kane, B.F.; Johnson, D.G.; Gorelick, R.J.; Henderson, L.E.; Levin, J.G. Zinc finger structures in the human immunodeficiency virus type 1 nucleocapsid protein facilitate efficient minus- and plus-strand transfer. Journal of virology 2000, 74, 8980–8988.

66. Thomas, J.A.; Gorelick, R.J. Nucleocapsid protein function in early infection processes. Virus research 2008, 134, 39–63, doi:10.1016/j.virusres.2007.12.006.

67. Rye-McCurdy, T.D.; Nadaraia-Hoke, S.; Gudleski-O’Regan, N.; Flanagan, J.M.; Parent, L.J.; Musier-Forsyth, K. Mechanistic differences between nucleic acid chaperone activities of the Gag proteins of Rous sarcoma virus and human immunodeficiency virus type 1 are attributed to the MA domain. J Virol 2014, 88, 7852–7861.

68. Stewart-Maynard, K.M.; Cruceanu, M.; Wang, F.; Vo, M.-N.; Gorelick, R.J.; Williams, M.C.; Rouzina, I.; Musier-Forsyth, K. Retroviral Nucleocapsid Proteins Display Nonequivalent Levels of Nucleic Acid Chaperone Activity. Journal of Virology 2008, 82, 10129, doi:10.1128/jvi.01169-08.

69. Chadwick, D.R.; Lever, A.M. Antisense RNA sequences targeting the 5’ leader packaging signal region of human immunodeficiency virus type-1 inhibits viral replication at post-transcriptional stages of the life cycle. Gene Ther 2000, 7, 1362–1368, doi:10.1038/sj.gt.3301254.

70. Ingemarsdotter, C.K.; Zeng, J.; Long, Z.; Lever, A.M.L.; Kenyon, J.C. An RNA-binding compound that stabilizes the HIV-1 gRNA packaging signal structure and specifically blocks HIV-1 RNA encapsidation. Retrovirology 2018, 15, 25, doi:10.1186/s12977-018-0407-4.

71. Andersson, A.-M.C.; Schwerdtfeger, M.; Holst, P.J. Virus-Like-Vaccines against HIV. Vaccines (Basel) 2018, 6, 10, doi:10.3390/vaccines6010010.

72. Zhao, C.; Ao, Z.; Yao, X. Current Advances in Virus-Like Particles as a Vaccination Approach against HIV Infection. Vaccines (Basel) 2016, 4, 2, doi:10.3390/vaccines4010002.

73. Doan, L.X.; Li, M.; Chen, C.; Yao, Q. Virus-like particles as HIV-1 vaccines. Reviews in Medical Virology 2005, 15, 75–88, doi:10.1002/rmv.449.

